# Development of a Thyroid Cancer to Lung Metastasis-on-a-Chip Model

**DOI:** 10.64898/2026.11.07.764982

**Authors:** Sydney R. Anderson, Tilak Khanal, Kylie G. Nairon, Aleksander Skardal, William E. Carson, Matthew D. Ringel

## Abstract

**Background:** Study of the metastatic cascade in differentiated non-medullary thyroid cancer (NMTC) is hampered by the relative lack of preclinical models that metastasize from primary tumors to other organs. We previously identified regulator of calcineurin 1 isoform 4 (RCAN1.4) as a suppressor of NMTC cell growth, invasion, and metastasis using *in vitro* systems and tail vein injections in immunocompromised mice. We then reported a microfluidic device mimicking the mouse tail vein injection where cancer cells were allowed to flow directly to lung organoid constructs. We now report the development of a full primary tumor to lung metastasis-on-a-chip (MOC) model for TC.

**Methods:** Resin molds for primary and secondary sites connected by a channel were 3D-printed and filled with polydimethylsiloxane to create a molded silicone elastomer. The primary site was seeded with an organoid composed of a hyaluronic acid and collagen-based hydrogel into which FTC236 and HTh74 NMTC cells with and without knockdown of RCAN1.4 were engrafted. The secondary site consisted of a hydrogel organoid into which bronchial epithelial cells and lung fibroblasts were seeded. Culture media was circulated and movement of NMTC cells to the secondary site was assessed.

**Results:** Both cell lines that expressed RCAN 1.4 metastasized to the secondary lung site. RCAN1.4-reduced FTC236 cells had increased metastatic behavior vs controls while HTh74 was less affected by RCAN1.4 loss, similar to prior studies. To determine if the MOC could be used for incorporation of immune components and therapeutic modeling, NK-92 cells were added and were shown to be reduce metastases for both cell lines in the presence or absence of RCAN1.4. Analysis of MOC circulating fluid revealed elevated levels of granzyme B, consistent with NK-92 cell activation. Cytokine profiling identified elevated probable NMTC and NK-92-derived factors.

**Conclusions:** These results indicate that NMTC metastatic behavior can be modeled using a novel MOC system for mechanistic and preclinical therapeutic studies.

## Introduction

Non-medullary thyroid cancer (NMTC) is the most common malignancy of primary endocrine organs. Patients with localized NMTC have an excellent prognosis following treatment with surgery and, in selected cases, radioiodine therapy and TSH suppression.^1^ In contrast, patients with progressive metastatic disease lack curative treatment options.^2^ Notably, patients with low volume metastatic well-differentiated NMTC can remain stable for decades. This characteristic makes NMTC an ideal model to identify mechanisms of metastatic dormancy.^3–6^ Genomic analysis of progressive metastatic lesions reveals that targetable new driver mutations are rare in the absence of dedifferentiation.^7,8^ Thus, we hypothesized that loss of metastasis progression suppressor genes may be relevant to progression of NMTC.^9^ These efforts led to identification of regulator of calcineurin 1.4 (RCAN1.4) as a new metastasis suppressor based on laboratory and clinical correlative studies.^9,10^ Clinical studies confirmed that the *RCAN1.4* promoter is frequently hypermethylated in NMTC leading to loss of expression. Functional experiments demonstrated activation of two transcription factors following the loss of RCAN1.4 in thyroid cells, NFATc2 and Nrf3 (encoded by *NFE2L3*).^9,11^ NFATc2 is negatively regulated by RCAN1.4 through direct competition for binding to the phosphatase calcineurin.^12^ NRF3 regulates thyroid cancer cell growth and invasiveness; and its overexpression is necessary for cancer progression following loss of RCAN1.4.^9,13^ Overexpression of *NFE2L3* induced more aggressive NMTC cell behavior *in vitro*, and in other cancers high levels of *NFE2L3* are associated with more aggressive features and a permissive immune environment.^9,13^ These data support the hypothesis that RCAN1.4 loss is functionally important in the NMTC metastasis.

To facilitate mechanistic studies of metastasis, our group developed poorly differentiated NMTC cells with stable reduction in RCAN1.4. These studies confirmed the growth and metastasis- suppression functions of RCAN1.4 in xenograft and tail vein injection models, but there are several limitations to these models. First, distant metastases rarely develop from NMTC xenografts. Second, if human NMTC cells are used, either immunocompromised or humanized mice are required. We have developed alternative methods to study NMTC metastases. We reported a metastasis-on-a-chip (MOC) system in which the human NMTC cells with or without RCAN1.4 knockdown were placed directly into the flow of a bioengineered construct mimicking a mouse tail vein model.^14^ NMTC cells with RCAN1.4 loss metastasized more quickly and were more invasive into the lung environment mirroring the murine studies. Consistent with prior work, RCAN1.4-reduced cells expressed elevated levels of NRF3 However, refinements of this metastasis model were needed to better replicate the metastatic process.

We now report development of an NMTC thyroid to lung cancer MOC in which cancer cells are placed in a primary tumor construct, enter flow through a narrow channel, and invade and grow in a second lung construct. Data from this model demonstrate that NMTC cells can metastasize from a primary site to a lung construct and that RCAN1.4-loss can enhance metastasis in some cells. Finally, we demonstrate that a natural killer cell line can reduce primary cancer construct growth and lung metastasis. This model may be useful for studying NMTC metastasis and for preclinical modeling.

## Materials and Methods

### Cell Lines and Cell Culture

Poorly differentiated NMTC cell lines with shRNA-induced stable knockdown of RCAN1.4 and control cell lines expressing a control shRNA developed by Wang, *et al*. were utilized.^9^ Their derivation from the parental FTC236 and HTh74 NMTC cell lines were previously reported.^9^ Briefly, the parental cell lines were the gifts of Dr. Rebecca Schweppe (University of Colorado, Denver, Colorado, USA) and their identities were confirmed by short tandem repeat profiling.^9,15^ Both cell lines express green fluorescent protein (GFP) and are cultured with puromycin to maintain knockdown.^9^ FTC236 cells are propagated in DMEM with 10% fetal bovine serum (FBS), 1% penicillin/streptomycin and 1% L-Glutamine. HTh74 cells are propagated in RPMI 1640 with 10% FBS and 1% non-essential amino acids. Cells are maintained in a humidified incubator at 37°C and 5% CO_2_.

## Microfluidic MOC fabrication

MOC microfluidic devices were fabricated using polydimethylsiloxane (PDMS) soft lithography.^14^ 3D printed molds were constructed from a computer assisted design (CAD) model that includes two orifices to serve as primary and secondary sites reflective of cancer cell invasion through a primary tumor site and metastasis towards a secondary location. The primary and secondary sites have an ellipsoid shape with a major axis of 6 mm and a minor axis of 5 mm. The primary site has a guide spot with a diameter of 2 mm. The secondary site is bisected into two semi circles by an open channel to allow for media flow to mimic vascular blood flow. Each MOC mold has four replicates of this primary to secondary site configuration arranged in parallel and connected via a linear channel for media flow (**Figure. 1**). Media is continuously perfused through the device over the 7-day experiment. Inlet and outlet ports are created using a 1 mm biopsy punch. The ports are placed directly below the primary site and directly above the secondary site. Flared PVC two-stop peristaltic pump tubing (0.51 mm internal diameter) (Perkin Elmer) is inserted into each port and connected to an Elemental Scientific MP2 microperistaltic pump. The tubing draws media from a 4 mL cryovial and is sealed with epoxy to prevent leakage. Cryovials have one hole for the outlet port tubing which is inserted 1.5 cm into the cryovial and another hole for the inlet port tubing which is inserted down to the bottom of the cryovial.

**Figure 1.**
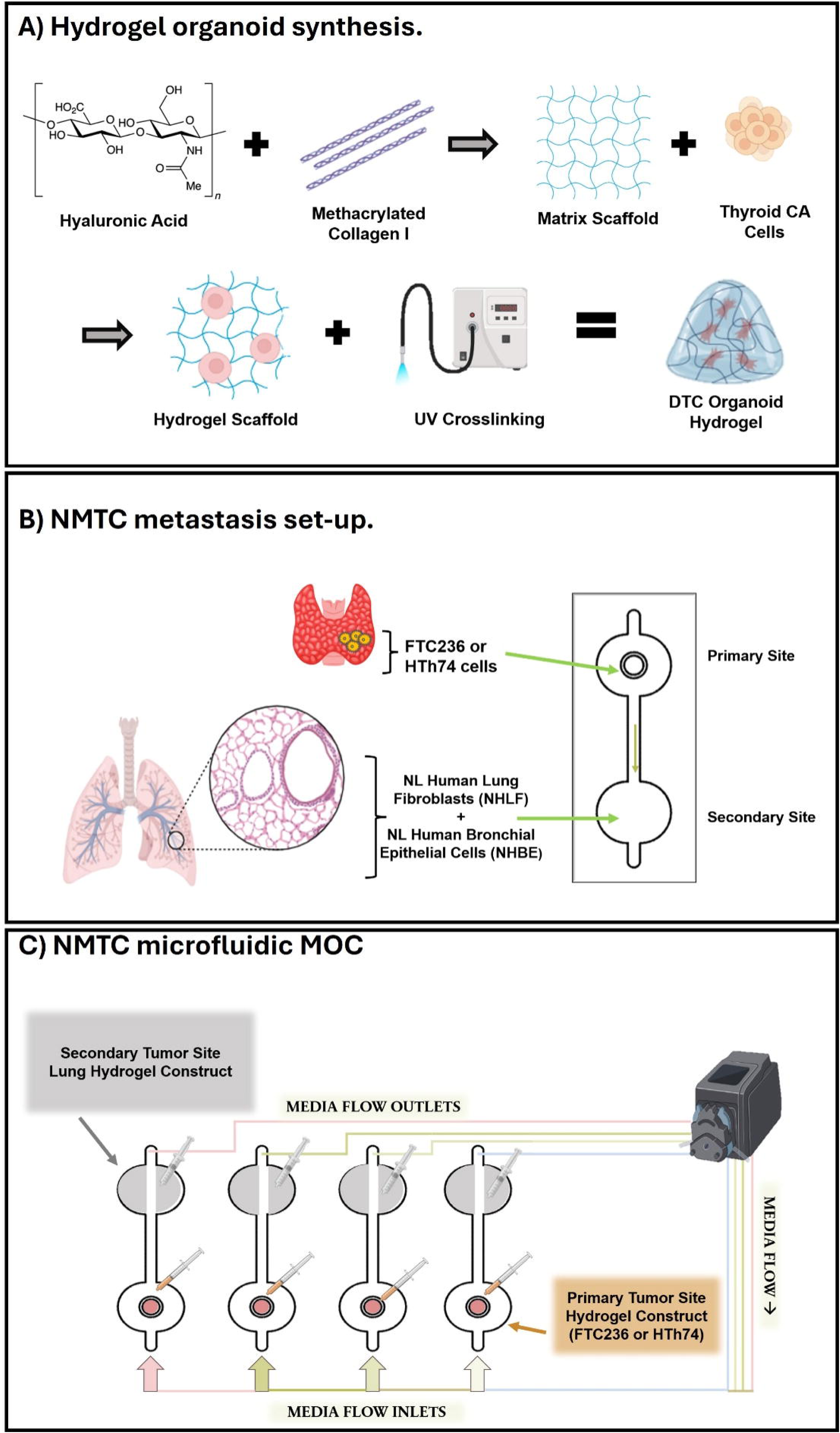
MOC design and experimental workflow. A) Hydrogel Organoid Synthesis. Hyaluronic acid, collagen I and NMTC cells are combined to give a hydrogel that is immobilized via UV cross-linking. B) Configuration of the primary and secondary sites within the MOC. C) Microfluidic channel flow patterns and placement of the pump device. Each channel is contained within an individual circuit. There are four circuits per MOC device. Therefore, there are 4 independent sets of tubing configurations (different colors in figure) that circulate around the microperistaltic pump.

Hollow molds were 3D printed with a digital light processing resin printer (H-Series, CadWorks 3D), sterilized using isopropyl alcohol, and cured in isopropyl alcohol baths followed by UV radiation for 20 minutes on each side. The molds were filled with PDMS using a 10:1 ratio of Sylgard 184 Elastomer:Curing Agent, sealed in a vacuum chamber to remove air bubbles, and cured overnight in a 65°C oven before the PDMS devices were removed.^16^ Constructs were sterilized with ethanol before being plasma bonded at 1 Torr to glass slides.

## Hydrogel encapsulation of cells and metastasis-on-a-chip operation

Hydrogel-based 3D tissue constructs are comprised of NMTC cells or lung cells (bronchial epithelial cells and lung fibroblasts) encapsulated within a collagen and hyaluronic acid-based hydrogel. Normal human bronchial epithelial (NHBE) cells and Normal human lung fibroblasts (NHLF) were purchased from Lonza, Inc. (Walkersville, MD).^17^ Methacrylated collagen I (PhotoColl, Advanced Biomatrix) was reconstituted at 6 mg/mL in 20 mM acetic acid on a rocking plate at 4℃ overnight. Thiolated hyaluronic acid (Heprasil, Advanced Biomatrix) was dissolved in 1X phosphate buffered saline solution at a pH of ∼7.4 and combined with a 10% photoinitiator solution (Irgacure) reaching a concentration of 1% w/v. NMTC cells were suspended at 10x10^6^ cells/mL in hydrogel precursor solution. NHBE and NHLF were suspended at a cell density of 5x10^6^ cells/mL in a 1:1 ratio. Cell-containing hydrogel constructs were photopolymerized using UV-light (BlueWave Redicure 365, Dymax) at 365 nm for 2 seconds following injection via insulin syringe at primary or secondary sites in the MOC devices.^14^ Consistent ∼10 µL hydrogel constructs were utilized across all experiments. A microperistaltic pump (Meinhard, Golden, CO) was used to maintain fluid flow at 47.4 µL/minute. A 1:1 ratio of fibroblast basal medium (FBM) complete with supplements (Lonza,) and bronchial epithelial cell growth medium (BEGM) with supplements (Lonza) was employed.

## NK-92 cell preparation

The NK-92 natural killer cell line (ATCC CRL-2407) was cultured in complete α-MEM media supplemented with 0.2 mM inositol, 0.1 mM 2-mercaptoethanol, 0.02 mM folic acid and 125 I.U./mL interleukin-2. NK-92 cells were fluorescently tagged using long-chain dialkylcarbocyanines (ThermoFisher) with an emission spectrum between 525-650 nm.^18,19^ For NK-92 experiments, the media reservoir contained 1:1:2 ratio of FBM complete, BEGM complete, and α-mem complete. supplemented with recombinant human IL-2 (Gibco, Peprotech).^20^ Media reservoirs contained 375,000 cells in 3 mL.

## Imaging

Images of the MOCs with or without NK-92 cells were taken on days 1, 4 and 7 after seeding. Imaging for MOCs containing NK-92 cells also included baseline images on day zero to confirm tumor cell loading prior to NK-92 cell infusion. A Nikon AXR Live Cell Imaging Confocal Microscope was used to obtain fluorescent and brightfield images of each hydrogel construct at both sites. Images were taken at 10x magnification in z-stacks of 200 µm with 21 steps.^14^

## Quantitation

Quantitative analyses of the Nikon AXR Confocal Images were conducted as previously described, with adjustment for cell density rather than migration distance, using IMAGEJ in which the images were uploaded into the software and background color and noise was removed.^14^ Images were thresholded and several measures were implemented to remove noise, debris, or artifact including watershed analysis and despeckling.^21^ Using the “analysis” feature, individual fluorescent objects were identified as cells and counted. Cell density counts were grouped by cell line and phenotype for data analysis.

## Enzyme Linked Immunosorbent Assay (ELISA)

Levels of granzyme B in circulating media were measured on day seven using the human granzyme B SimpleStep ELISA Kits (ab212163 and ab235635, Abcam) according to manufacturer instructions. Absorbance was measured at 450 nm using a plate reader (VarioSkan Lux, ThermoFisher).

## Cytokine Analysis

Cytokine analysis of the circulating media was performed by collecting duplicate samples of each experimental condition on day seven. Samples were shipped to RayBioTech for analysis of targets (Human Cytokine Array Q4000, QAH-CAA-4000-1, RayBiotech, Inc., Norcross, GA).^22^

## Statistical Analysis

All statistical analyses were conducted using Microsoft Excel or GraphPad/Prism. All experiments were performed with at least two biological replicates. To determine statistical significance between experimental groups, a t-test or a one-way analysis of variance (ANOVA) followed by Tukey’s multiple comparison test was conducted. A p-value ≤0.05 was considered statistically significant.

## Results

### Design of a 3D MOC model and experimental workflow

Building upon the prior model in which tumor cells flowed directly to a lung metastatic secondary site, the present model contains a primary site consisting of a hyaluronic acid and collagen- based hydrogel into which TC cells were seeded, and the secondary site consists of this hydrogel into which NHBE and NHLF were seeded.^14^ Continuous flow of circulating media between the two sites was established using a microperistaltic pump. Tumor cells were required to “intravasate” into the circulation and then “extravasate” into the lung hydrogel and survive. A schematic and photo of the MOC device and overall experimental workflow are provided in **Figure 1**.

## NMTC cells metastasize in the MOC system

This metastasis model was used to determine if the NMTC cells could metastasize in the MOC using the validated RCAN1.4 knockdown and control FTC236 and HTh74 cells from prior studies.^9,14,16,23^ Following initiation of the MOC devices on day 0, confocal microscopy images overlaid onto brightfield images were used to visualize the presence of fluorescently labeled NMTC cells in both the primary and lung constructs on days 1, 4, and 7. For both the FTC236 and HTh74 cell lines, evidence of cell movement from the primary tumor to the lung construct was identified for the scramble controls, confirming the ability of the cells to metastasize in the MOC. The FTC236 G9 and F4 RCAN1.4 and the HTh74 knockdown cells demonstrated similar or increased metastasis versus scramble controls (**Figures 2A and 2B)**.

**Figure 2:**
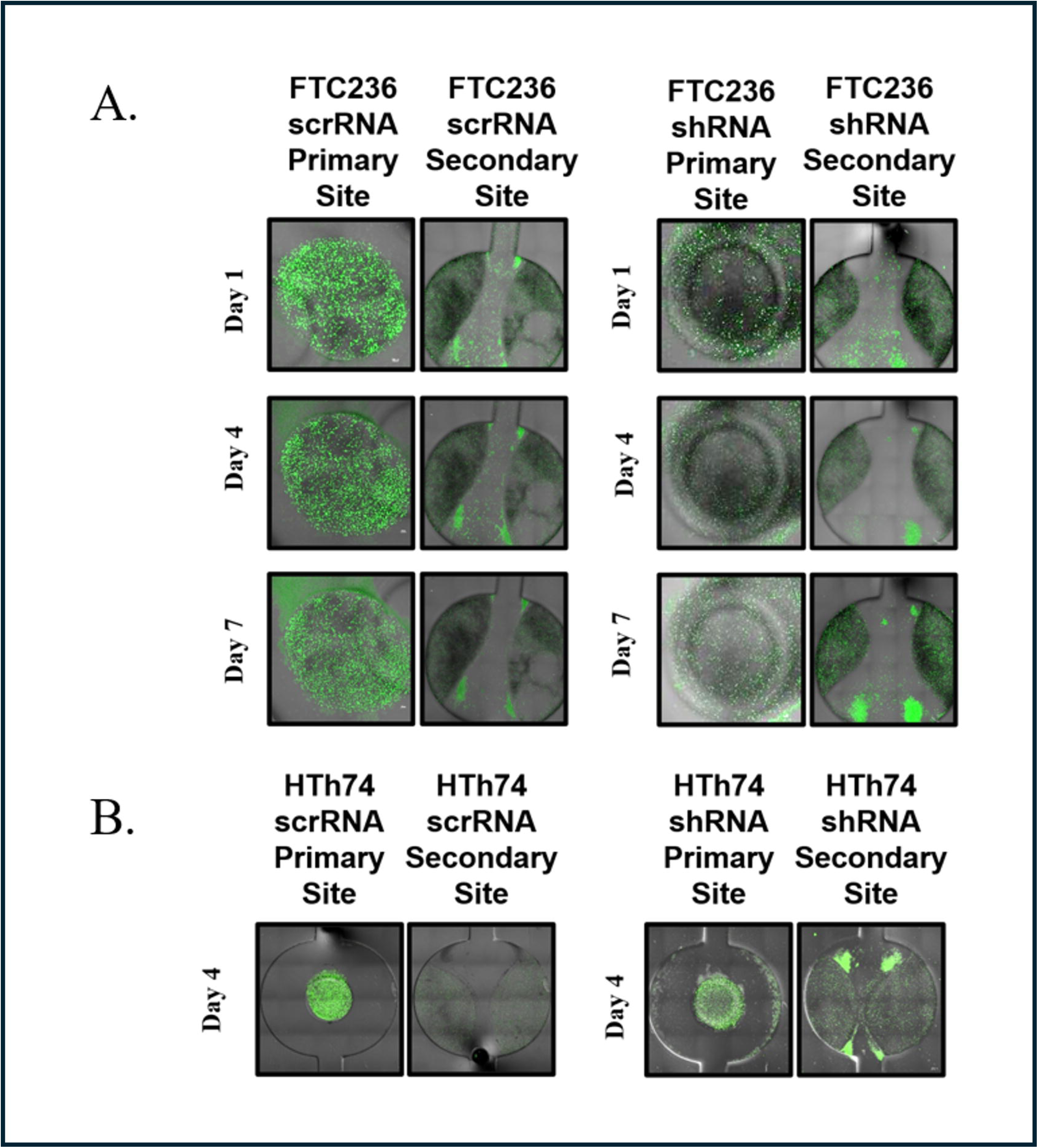
**NMTC cell movement from primary tumor to pulmonary site in the MOC system**. Fluorescent microscopy images of FTC236 TC cells within the primary tumor and secondary pulmonary sites of four MOC devices arrayed in parallel. A). FTC236 cells treated with control (scrRNA) or RCAN1.4 knockdown (shRNA) constructs were seeded at the primary site and the MOC was imaged on days 1, 4, and 7. Representative images of connected primary and secondary sites are shown. NMTC cells fluoresce green. B). Images obtained from a similar experiment that employed HTh74 TC cells treated with scrRNA or RCAN1.4 shRNA, day four images.

To quantify the MOC images, an image binarization approach was employed.^14,24,25^ Fluorescent images were converted to black and white to facilitate quantitation of cell densities. All clones populated the lung lesions and using this approach; one FTC236 RCAN1.4 knockdown more effectively populated the lung constructs compared to scramble control (**Figure 3A**). HTh74 RCAN1.4 knockdown cells demonstrated increased cell density over time and were similar to the control cell line metastases by Day 7 (**Figure 3A)**. Using these images, the number of metastatic NMTC cells that had invaded the lung constructs were quantified. Metastasis was seen for all cell lines but was increased with RCAN1.4 knockdown, albeit with different kinetics for each cell line (**Figure 3B)**. On day 4, there was a significant 6.5-fold increase in FTC236 G9 clone cells at the metastatic site (clone 2; p= 0.041) and a non-significant 1.5-fold increase in the F4 clone (clone 1). In HTh74 cells there was a non-significant 1.5-fold increase in cells at the lung site by day seven but not at day four. Notably, quantifiable metastasis to the lung constructs was identified for both cell lines regardless of RCAN1.4 status providing proof-of- principle for the MOC.

**Figure 3:**
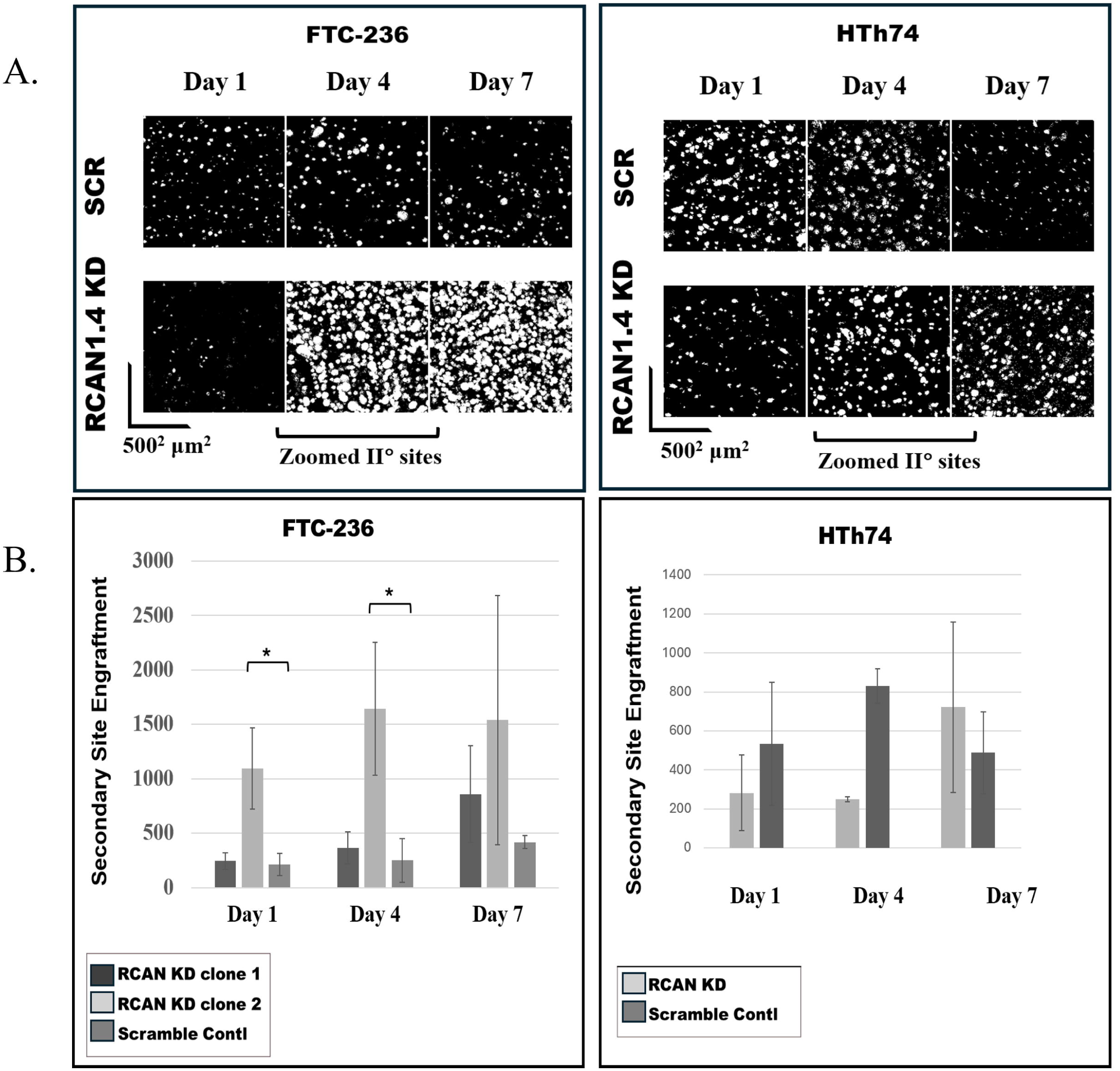
Quantitation of primary-to-lung NMTC metastases. A) Thresholded images (left) of FTC236 scrRNA and RCAN1.4 knockdown cells (clone F4) at 1, 4, and 7 days. Images (right) for HTh74 cell line at those same timepoints. B). Bar graph depiction of FTC236 secondary site engraftment (left). Results (right) for the HTh74 cell line. Statistics were conducted using Student’s t test and multiple unpaired t tests with * p < 0.05, ** p < 0.01, *** p < 0.005, and **** p < 0.0005. Error bars represent standard deviation (SD).

## NK cells inhibit NMTC cell metastasis in the MOC model

There is expanding interest in NK cell therapy for NMTC.^26^ CAR NK cells have shown benefit in murine NMTC models.^27,28^ To determine suitability of the MOC model for preclinical studies, the NK-92 natural killer cell line was integrated into the system. MOC devices were initiated, fluorescently stained NK-92 cells were infused into the circulating fluid at the outset of the experiments and were observed in the primary and secondary sites at all time points. In the presence of NK-92 cells, RCAN1.4-knockdown and control FTC236 and HTh74 cells were reduced at the primary site by day 7 (**Figure 4A, 4B**). In addition, in comparison to MOC without NK-92 cells (**Figure 2**), the NK-92 cells markedly reduced thyroid cell movement to the secondary site for both the scramble control and RCAN1.4 knockdown NMTC cells. (**Figure 4C**). Image thresholding enabled quantification revealed a reduction in cell numbers at the secondary site with NK-92 infusion (**Figure 4D** **vs** **Figure 3**).

**Figure 4:**
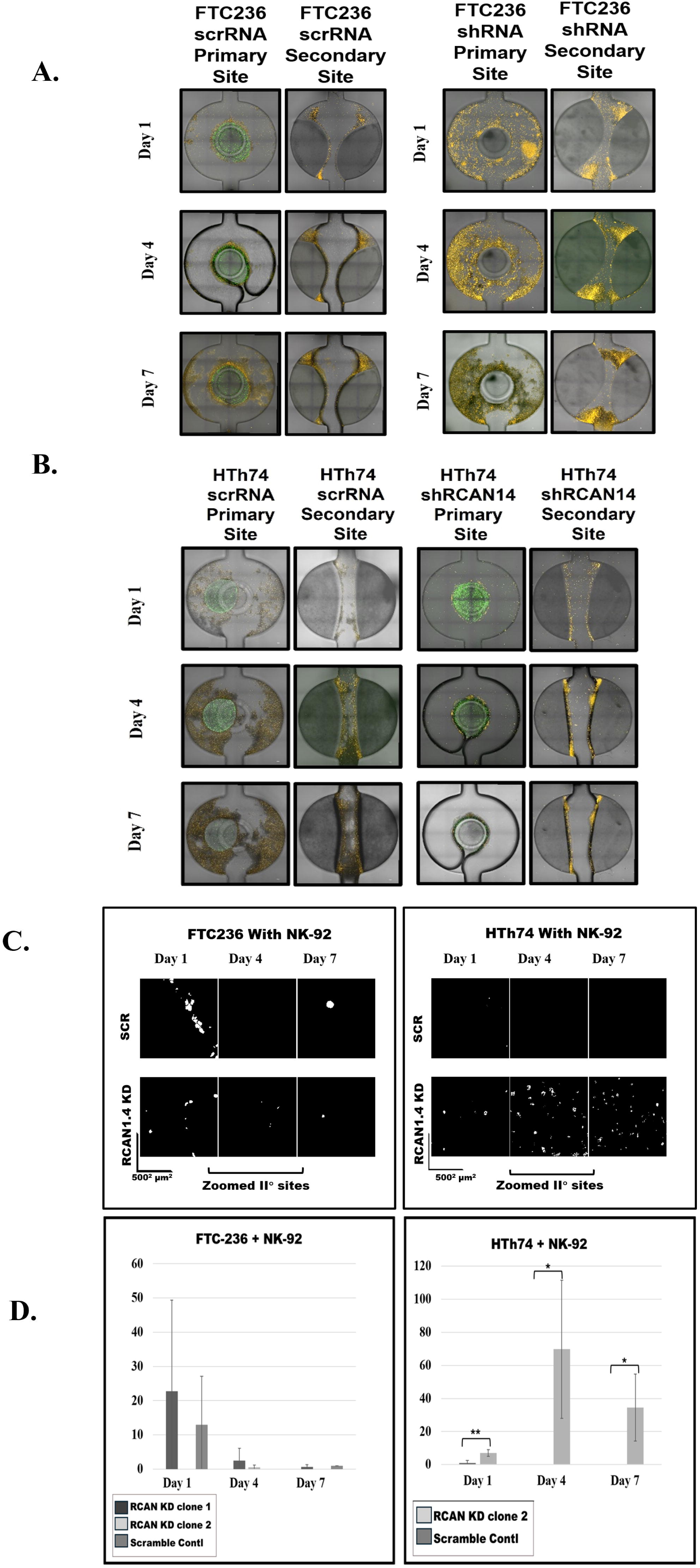
Addition of NK-92 cells to the MOC. Fluorescent microscopy images of FTC236 NMTC cells within the primary tumor and secondary pulmonary sites of four MOC devices arrayed in parallel. Fluorescently labeled NK-92 cells added to the systems on day zero and appear orange. A). FTC236 cells expressing control (scrRNA) or RCAN1.4 knockdown (shRNA) were seeded at the primary site and the MOC was imaged on days 1, 4, and 7. Representative images of connected primary and secondary sites are shown. B). Images obtained from a similar experiment that employed HTh74 NMTC cells treated with scrRNA or RCAN1.4 shRNA. C). Thresholded images of the devices presented in A and B compared to those obtained in Figure 2. D). Bar graph depiction of the results of the NK-92 experiment compared to those obtained in the absence of NK-92 cells. Statistics were conducted using Student’s t test and multiple unpaired t tests with * p < 0.05, ** p < 0.01, *** p < 0.005, and **** p < 0.0005. Error bars represent standard deviation (SD).

## NK-92 cells aggregate around NMTC cells and release granzyme B

Confocal microscopy images of the MOC on Day 0 suggested aggregation of NK cells around control and RCAN1.4 KD NMTC cells at the primary site (**Figure 5A, B**). Granzyme B is released by activated NK cells and its levels in day 7 MOC media were assayed. Increased levels were identified in MOC systems containing NK-92 cells and addition of NK-92 cells to the RCAN1.4 knockdown system led to significantly higher levels of granzyme B compared to the scramble control (**Figure 5C**).

**Figure 5:**
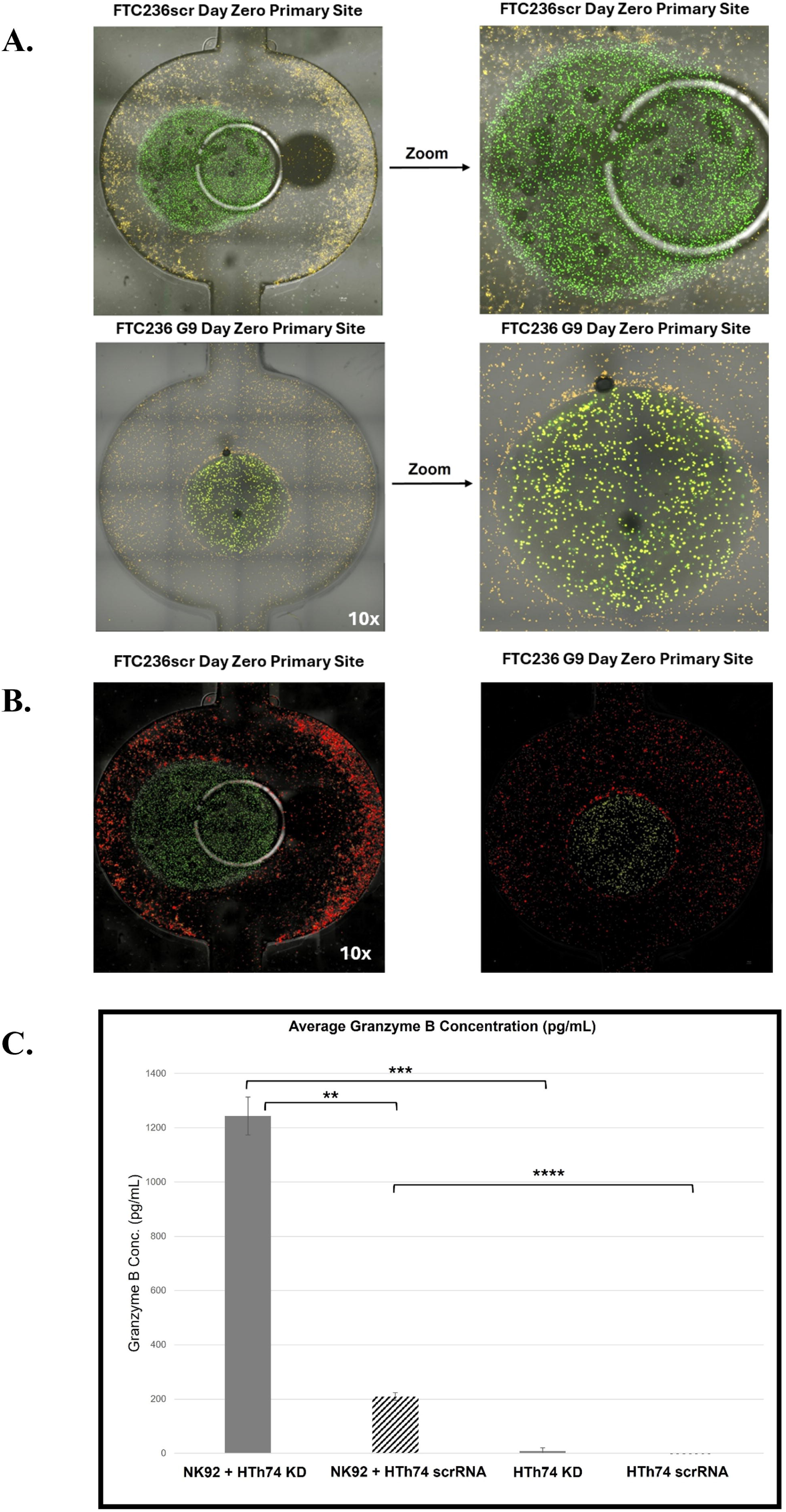
NK cells release granzyme B when placed into the MOC with NMTC cells. A). Day zero images of a representative primary site (FTC236 scrRNA and G9 knockdown) demonstrating the presence of NMTC cells and NK-92 cells with a localization “halo” around the primary construct: 10x and zoomed images. B). Color threshold based on dye color. Orange (NK cell) and green (thyroid cells) filtered using the Hue function in IMAGEJ for FTC236 scramble (left) and G9 (right). C) Levels of granzyme B with the circulating media were measured by ELISA on day seven for HTh74 cells. Statistics were conducted using Student’s t test with * p < 0.05, ** p < 0.01, *** p < 0.005, and **** p < 0.0005. Error bars represent standard deviation (SD).

## Cytokine profile of NK-92 MOC

To screen for other potential secreted factors in the MOC, culture supernatants were analyzed to determine the cytokine profiles released into the NMTC MOC supernatant by the cancer cells alone **(Fig. 6A**) or following addition of the NK-92 cells (**Fig 6B**). In the absence of NK-92 cells, GDF-15, IL-8, TIMP-1 and MIF were elevated in FTC236 RCAN1.4 knockdown cells and TIMP- 2 was increased in HTh74 RCAN1.4 knockdown cells versus their scramble controls (**Fig. 6A**). In the presence of NK-92 cells, seventeen cytokines were elevated in the supernatants from MOCs including NK cell-derived factors (e.g., Flt-3 ligand, Fas ligand, MIP-1α, MIP-1β, RANTES, TIMP-2) and presumed NMTC cell-secreted factors such as heparin binding epithelial growth factor (HB-EGF), insulin-like growth factor binding protein 2 (IGFBP-2), epidermal growth factor receptor (EGFR), osteoprotegerin (OPG) and TNF-related apoptosis-inducing ligand receptor 3 (TRAIL-R3) (**Fig. 6B**).

**Figure 6.**
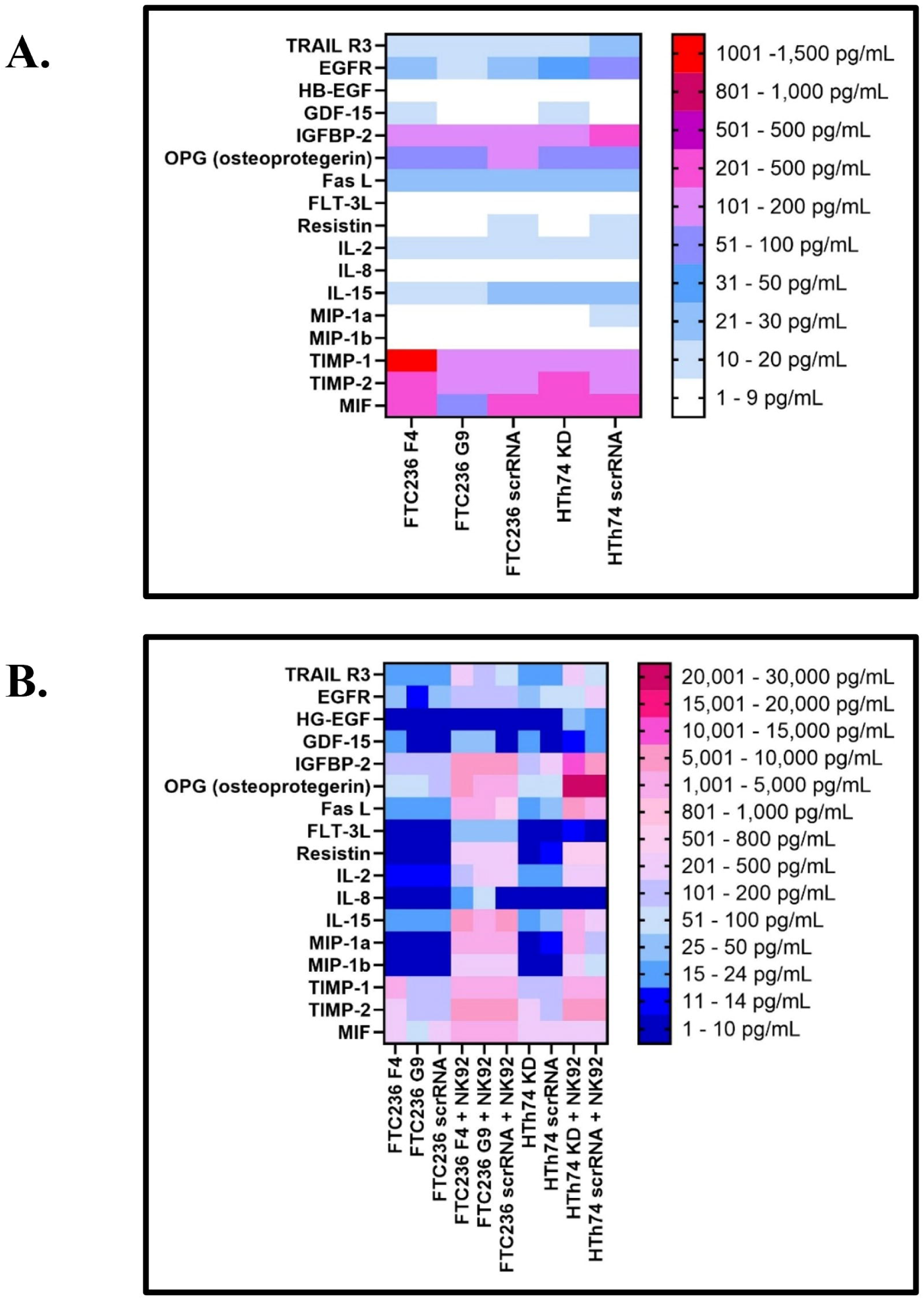
Altered cytokine levels in response to RCAN1.4 knockdown and addition of NK-92 cells. Day 7 culture media was assayed for levels of 200 cytokines by a multiplexed assay. Results are presented as heat maps. A). Loss of RCAN1.4 expression caused upregulation of GDF-15, IL-8, MIP-1α, and TIMP-1. B) Addition of NK-92 cells into circulating media led to increased levels of factors thought to be derived from NK-92 cells (Flt-3 ligand, Fas ligand, MIP- 1α, MIP-1β, RANTES, TIMP-2, MIF) and factors thought to be the product of NMTC cells such TNF-Related apoptosis-inducing ligand receptor 3 (TRAIL-R3), epidermal growth factor receptor (EGFR), heparin binding epithelial growth factor (HB-EGF), insulin-like growth factor binding protein 2 (IGFBP-2). IL-2 was detected due to its inclusion in the culture media.

## Discussion

The present study to our knowledge provides the first description of a NMTC primary tumor to lung metastasis model utilizing a MOC platform. The metastatic capabilities of two NMTC cell lines with and without RCAN1.4 knockdown were similar to prior studies in a simplified MOC and in a murine xenograft model.^9,14^ In the present model, cancer cells must invade out of a hydrogel primary site, travel through a model circulatory system and invade a second hydrogel construct partially mimicking the lung environment. This new MOC model is an important step in filling a critical gap in NMTC modeling and has the potential to facilitate mechanistic and preclinical studies.

For the present studies, human NMTC cell lines with baseline and shRNA-reduced levels of RCAN1.4 were employed.^3^ RCAN1.4 is hypermethylated and silenced in human TC samples, and its loss is associated with metastases.^3,29^ In the present study, both poorly differentiated NMTC cell lines metastasized in the MOC. The FTC236 cell line exhibited a more rapid and more extensive movement to the secondary site when RCAN1.4 was reduced while the impact of RCAN1.4 loss was less obvious in the HTh74 cell line. These results are similar to our prior reports using tail vein and direct flow models with these cell lines.^9,14^ Further studies with additional cell lines, including patient-derived NMTC samples, and in models of increasing biological complexity will be required to fully elucidate the mechanisms underlying the effects of RCAN1.4 loss on metastatic progression.

Potential applications of this system include mechanistic studies of cancer progression in different local environments, and the ability to characterize inhibitors of different phases of the metastatic process. As a proof-of-principle, the IL-2-dependent NK-92 cell line was incorporated into this flow-based model.^27^ This cell line has been employed elsewhere in clinical trials for human cancer, and modified versions have been created to target specific cell surface antigens such as the thyroid-stimulating hormone (TSH) on NMTC cells.^30^ Day zero images reveal that NK-92 cells interacted with NMTC cells and released granzyme B, consistent with activation. NK-92 cells appeared to prevent metastatic engraftment, including cells with reduced RCAN1.4 levels. Future studies will determine if targeted therapies directed at thyroid-cancer cells are also effective in such a system.

There are several weaknesses of the current system. While the current primary tumor location includes NMTC cells in a complex hydrogel matrix; additional cell types will be required for more accurate modeling. Also, the blood vessel model does not include cellular components of vessel walls, such as endothelial cells and pericytes. These components are being optimized to better model intravasation, circulation, and extravasation. While the lung site contains multiple normal lung cell-types, it does not yet fully reflect lung structure or oxygenation saturation which may be important in tumor growth and response. NK-92 cells were employed, and they have been used in clinical studies; however, use of CAR-NK, CAR-T and other therapies are being explored. Finally, the use of patient-derived tumors and their tumor-associated immune cells is being explored to improve preclinical modeling.

In summary, a primary-to-metastatic NMTC MOC system was developed using two control NMTC cell lines and their RCAN1.4-reduced clones. All cell lines metastasized from the thyroid site to the lung construct over one week with evidence of regulation by RCAN1.4. The human NK-92 cell line was integrated into this model and inhibited metastatic progression that coincided with activation. These data provide evidence of feasibility of this MOC model for NMTC.

## Acknowledgements

The authors gratefully acknowledge funding by the Department of Defense W81XWH-22-1-0647 (MDR and AS), R01 CA240302 (MDR), and support from NIH grant P30 CA016058. Special thanks to Sandya, Liyanarachchi, MS for assistance with the statistical analyses.

## References

1 Ringel, M. D. et al. 2025 American Thyroid Association Management Guidelines for Adult Patients with Differentiated Thyroid Cancer. Thyroid 35, 841–985 (2025). 10.1177/10507256251363120

2 Nguyen, Q. T. et al. Diagnosis and treatment of patients with thyroid cancer. Am Health Drug Benefits 8, 30–40 (2015).

3 Rajan, N., Khanal, T. & Ringel, M. D. Progression and dormancy in metastatic thyroid cancer: concepts and clinical implications. Endocrine 70, 24–35 (2020). 10.1007/s12020-020-02453-8

4 Valenciaga, A., Ivancic, R. J., Khawaja, R., Way, D. P. & Bahner, D. P. Efficacy of an Integrated Hands-On Thyroid Ultrasound Session for Medical Student Education. Cureus 13, e12421 (2021). 10.7759/cureus.12421

5 Chan, C. M. et al. Targeted inhibition of Src kinase with dasatinib blocks thyroid cancer growth and metastasis. Clin Cancer Res 18, 3580–3591 (2012). 10.1158/1078-0432.Ccr-11-3359

6 Ma, Y. et al. Development of p21 activated kinase-targeted multikinase inhibitors that inhibit thyroid cancer cell migration. J Clin Endocrinol Metab 98, E1314–1322 (2013). 10.1210/jc.2012-3937

7 Justiniano, S. E. et al. Genetic variants in thyroid cancer distant metastases. Endocr Relat Cancer 23, L33–36 (2016). 10.1530/erc-16-0351

8 Sohn, S. Y. et al. Highly Concordant Key Genetic Alterations in Primary Tumors and Matched Distant Metastases in Differentiated Thyroid Cancer. Thyroid 26, 672–682 (2016). 10.1089/thy.2015.0527

9 Wang, C. et al. RCAN1-4 is a thyroid cancer growth and metastasis suppressor. JCI Insight 2, e90651 (2017). 10.1172/jci.insight.90651

10 Stathatos, N. et al. KiSS-1/G protein-coupled receptor 54 metastasis suppressor pathway increases myocyte-enriched calcineurin interacting protein 1 expression and chronically inhibits calcineurin activity. J Clin Endocrinol Metab 90, 5432–5440 (2005). 10.1210/jc.2005-0963

11 Espinosa, A. V. et al. Regulator of calcineurin 1 modulates cancer cell migration in vitro. Clin Exp Metastasis 26, 517–526 (2009). 10.1007/s10585-009-9251-1

12 Rothermel, B. et al. A Protein Encoded within the Down Syndrome Critical Region Is Enriched in Striated Muscles and Inhibits Calcineurin Signaling *. Journal of Biological Chemistry 275, 8719–8725 (2000). 10.1074/jbc.275.12.8719

13 Kilicaslan, D., Irmak-Yazicioglu, M. B. & Copuroglu, E. Nrf3: an emerging player in cancer, inflammation, and cellular homeostasis. Mol Biol Rep 53 (2026). 10.1007/s11033-026-11678-1

14 Nairon, K. et al. RCAN1.4 regulates tumor cell engraftment and invasion in a thyroid cancer to lung metastasis-on-a-chip microphysiological system. Biofabrication (2024). 10.1088/1758-5090/ad82e0

15 Schweppe, R. E. et al. Deoxyribonucleic acid profiling analysis of 40 human thyroid cancer cell lines reveals cross-contamination resulting in cell line redundancy and misidentification. J Clin Endocrinol Metab 93, 4331–4341 (2008). 10.1210/jc.2008-1102

16 Dedhia, P. H. et al. A 3D adrenocortical carcinoma tumor platform for preclinical modeling of drug response and matrix metalloproteinase activity. Scientific Reports 13, 15508 (2023). 10.1038/s41598-023-42659-0

17 Mazzocchi, A., Devarasetty, M., Huntwork, R., Soker, S. & Skardal, A. Optimization of collagen type I-hyaluronan hybrid bioink for 3D bioprinted liver microenvironments. Biofabrication 11, 015003 (2018). 10.1088/1758-5090/aae543

18 Probes, M. Vybrant™ Cell-Labeling Solutions.

19 Invitrogen. Lipophilic Tracers—Dil, DiO, DiD, DiA, and DiR.

20 Rodriguez, M. A. et al. Establishment of a patient-derived adrenocortical carcinoma 3D tumor construct platform for evaluation of therapeutic strategies. bioRxiv (2026). 10.64898/2026.01.08.698468

21 ImageJ User Guide Revised edition. 1.46r edn, 34, 64-75, 116, 130-132, (2012).

22 Nairon, K. G., DePalma, T. J., Zent, J. M., Leight, J. L. & Skardal, A. Tumor cell-conditioned media drives collagen remodeling via fibroblast and pericyte activation in an in vitro premetastatic niche model. iScience 25, 104645 (2022). 10.1016/j.isci.2022.104645

23 Skardal, A., Sivakumar, H., Rodriguez, M. A., Popova, L. V. & Dedhia, P. H. Bioengineered in vitro three-dimensional tumor models in endocrine cancers. Endocr Relat Cancer 31 (2024). 10.1530/erc-23-0344

24 Aleman, J. & Skardal, A. A multi-site metastasis-on-a-chip microphysiological system for assessing metastatic preference of cancer cells. Biotechnol Bioeng 116, 936–944 (2019). 10.1002/bit.26871

25 Aleman, J. et al. Deconstructed Microfluidic Bone Marrow On-A-Chip to Study Normal and Malignant Hemopoietic Cell-Niche Interactions. Small 15, e1902971 (2019). 10.1002/smll.201902971

26 Seok, J. et al. Tumor microenvironment remodeling across thyroid cancer differentiation states revealed by spatial transcriptomics. Cancer Immunology, Immunotherapy 74, 357 (2025). 10.1007/s00262-025-04210-0

27 Klingemann, H. The Natural Killer Cell Line NK-92 and Its Genetic Variants: Impact on NK Cell Research and Cancer Immunotherapy. Cancers (Basel) 17 (2025). 10.3390/cancers17121968

28 Zhou, J. et al. Development of TSHR-CAR NK-92 cells for Differentiated Thyroid Cancer. Molecular and Cellular Endocrinology 589, 112251 (2024). 10.1016/j.mce.2024.112251

29 Khanal, T., Rajan, N., Li, W., Liyanarachchi, S. & Ringel, M. D. The RCAN1.4 Metastasis Suppressor Is Hypermethylated at Intron 1 in Thyroid Cancer. Thyroid 33, 965–973 (2023). 10.1089/thy.2022.0687

30 Zhou, J. et al. Development of TSHR-CAR NK-92 cells for Differentiated Thyroid Cancer. Mol Cell Endocrinol 589, 112251 (2024). 10.1016/j.mce.2024.112251

